# Role of hypothalamic MAPK/ERK signaling in diabetes remission induced by the central action of fibroblast growth factor 1

**DOI:** 10.1101/2020.12.24.424313

**Authors:** Jenny M. Brown, Marie A. Bentsen, Dylan M. Rausch, Bao Anh Phan, Danielle Wieck, Huzaifa Wasanwala, Miles E. Matsen, Nikhil Acharya, Nicole E. Richardson, Xin Zhao, Peng Zhai, Anna Secher, Gregory J. Morton, Tune H. Pers, Michael W. Schwartz, Jarrad M. Scarlett

**Author notes:** Address for Correspondence: Jarrad M. Scarlett, University of Washington Medicine Diabetes Institute, Department of Medicine, 750 Republican St, F770, Seattle, Washington, 98195, USA.

## Abstract

The capacity of the brain to elicit sustained remission of hyperglycemia in rodent models of type 2 diabetes following intracerebroventricular (icv) injection of fibroblast growth factor 1 (FGF1) is well established. Here, we show that following icv FGF1 injection, hypothalamic signaling by extracellular signal-regulated kinases 1 and 2 (ERK1/2), members of the mitogen-activated protein kinase (MAPK) family is induced for at least 24h. Further, we show that in diabetic Lep^*ob/ob*^ mice, this prolonged response is required for the sustained antidiabetic action of FGF1, since it is abolished by sustained (but not acute) pharmacologic blockade of hypothalamic MAPK/ERK signaling. We also demonstrate that FGF1 R50E, a FGF1 mutant that activates FGF receptors but induces only transient hypothalamic MAPK/ERK signaling, fails to mimic the sustained glucose lowering induced by FGF1. These data identify sustained activation of hypothalamic MAPK/ERK signaling as playing an essential role in the mechanism underlying diabetes remission induced by icv FGF1 administration.

## Introduction

Fibroblast growth factor 1 (FGF1) is a prototypical FGF that, in addition to its role in biologic functions ranging from brain development to angiogenesis and wound repair, is implicated the in regulation of feeding and glucose homeostasis^1,2^. Recently, FGF1 emerged as a potential antidiabetic agent that following a single intracerebroventricular (icv) injection, induces remission of diabetic hyperglycemia lasting weeks or months in both mouse (Lep^*ob/ob*^ and LepR^*db/db*^) and rat (Zucker Diabetic Fatty (ZDF)) models of type-2 diabetes (T2D)^3-6^. Since the effect of icv FGF1 administration to normalize diabetic hyperglycemia in a sustained manner is recapitulated by microinjection of a lower dose of FGF1 directly into mediobasal hypothalamus (MBH), this brain area is implicated as a key target for this sustained antidiabetic response^5^. While recent work points to a role for glia-neuron interactions leading to increased melanocortin signaling^7^, as well as to FGF1-induced changes in MBH extracellular matrix^8^, the signal transduction mechanisms by which FGF1 action in the MBH induces sustained remission of diabetic hyperglycemia is unknown.

Among various intracellular signaling cascades activated by binding of FGF1 to FGF receptors (FGFRs) is the extracellular-signal-regulated kinase (ERK) pathway, a major signaling cassette of the mitogen activated protein kinase (MAPK) signal transduction system that communicates signals arising from activation of cell surface receptors to changes of gene expression occurring in the cell nucleus^9^. Previous work demonstrates that MAPK/ERK signaling is rapidly induced in the hypothalamus of diabetic mice by FGF19, a member of the endocrine FGF family, and that this ERK activation is required for its transient glucose-lowering action^10^. Activation of ERK signaling in the MBH has also been documented following icv administration of FGF1 and is associated with robust transcriptional changes related to the ERK pathway in both tanycytes and astrocytes^5,7^. However, neither the role played by ERK signaling in these responses nor the extent to which they contribute to the sustained antidiabetic effect of icv FGF1 is known. The current work was undertaken to address these questions.

We report that in diabetic Lep^*ob/ob*^ mice, the sustained antidiabetic effect of icv FGF1 is abolished by pharmacologic blockade of hypothalamic MAPK/ERK signaling (achieved by central administration of the MAPK inhibitor U0126), but only if this blockade lasts for the 24h duration of FGF1-induced activation of MAPK/ERK signaling in the hypothalamus. We further report that icv injection of FGF1 R50E, a FGF1 mutant that activates FGFRs and induces transient, but not sustained MAPK/ERK signaling^11,12^, fails to mimic the sustained diabetes remission induced by icv FGF1 administration. We also report transcriptomic evidence pointing to a role for astrocytes (and possibly tanycytes) as primary targets for the hypothalamic response to FGF1. Collectively, these data indicate that durable induction of hypothalamic MAPK/ERK signaling is required for sustained remission of diabetic hyperglycemia induced by icv FGF1 administration and adds to growing evidence of a role for glia-neuron interaction in this response.

## Results

### Central Administration of FGF1 Induces Sustained ERK1/2 Signaling in the Hypothalamus

Consistent with our previous observations in ZDF rats^5^, we found using quantitative western blot that in overnight-fasted wild-type (WT) mice, phosphorylation of ERK1/2 (pERK1/2, a biomarker of ERK1/2 activation) was detectable in the hypothalamus within 20 min of icv injection of FGF1 (3 µg), but not following icv vehicle injection (Figure 1A). Since the duration of ERK1/2 activation is a determinant of gene expression and other cellular processes engaged by ERK signaling^13^, we next measured the time course of ERK1/2 activation in the hypothalamus following a single icv injection of FGF1. We found that in WT mice, activation of ERK1/2 was readily detected at both the 5h (Figure 1B) and 24h (Figure 1C) time points following icv FGF1 injection, but this response was no longer evident at 48h (Figure 1D). In contrast, changes of hypothalamic pERK1/2 content were not observed at any time point after icv vehicle. Interestingly, the magnitude of FGF1-induction of ERK1/2 increased over time, being nearly 3-fold greater at 24h than at 20 min.

**Figure 1.**
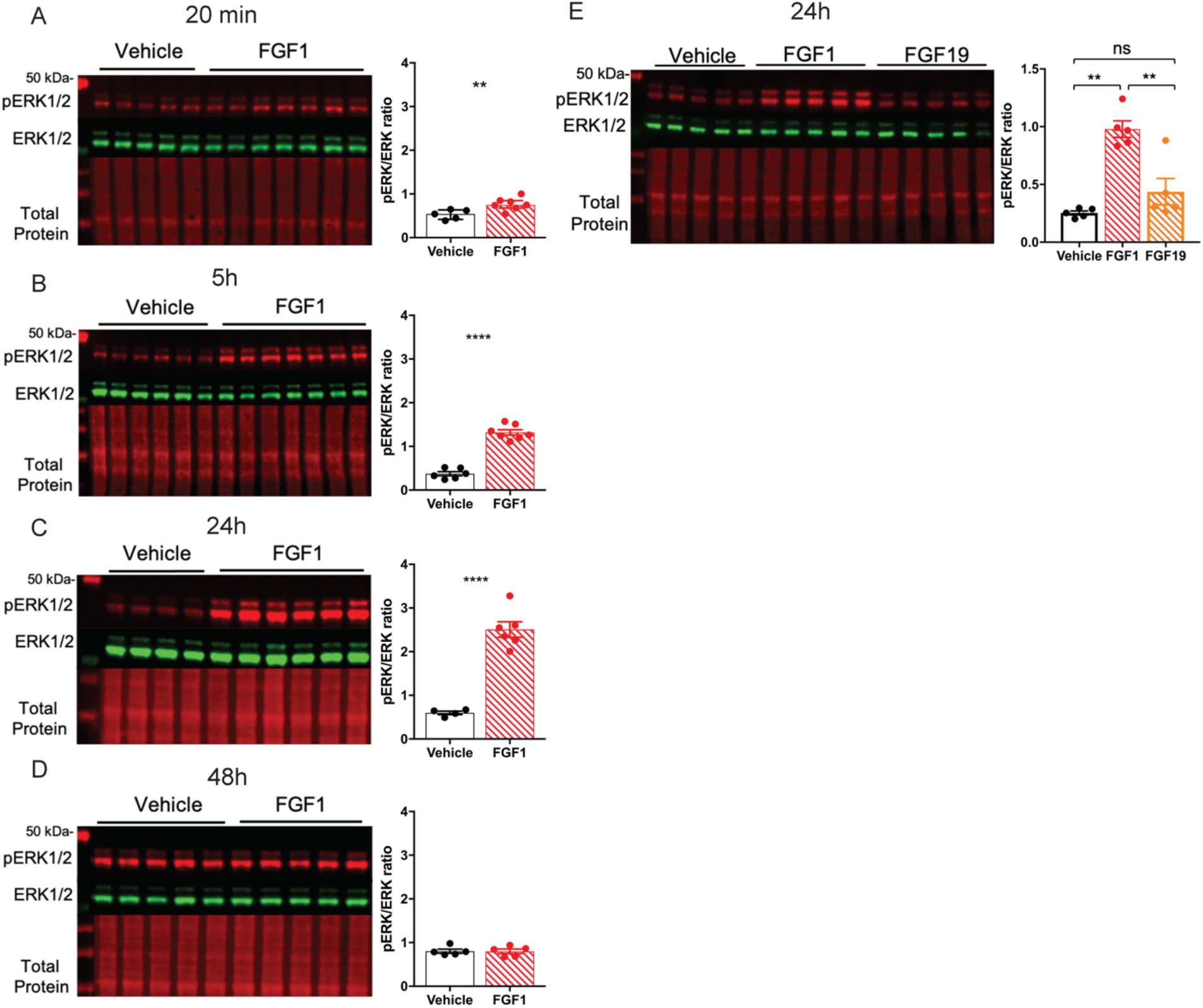
Central Administration of FGF1 Induces Sustained ERK1/2 Signaling in the Hypothalamus. Representative western blot (left panel) and quantitative comparison (right panel) of phosphorylated (red) and total ERK1/2 (green) and total protein (red) from hypothalamic of punches from adult male C57Bl6J mice after a single icv injection of either vehicle or FGF1 (3 μg) at **A)** 20 minutes (n=5-7/group, t=3.2497 df=9.8554 p=0.00444) **B)** 5 hours (n=6-7/group, t=11.937 df=10.624 p=8.62e-08) **C)** 24 hours (n=4-6/group, t=10.528, df=5.4845, p=3.816e-05) and **D)** 48 hours (n=5/group, t=-0.0344, df=7.8827, p=0.5133) after injection. pERK1/2 ratio unaired Welch two sample t-test (one sided). **P<0.01, ****P<0.0001 **E)** Quantitative western blot of ERK1/2 phosphorylation 24h after a single icv injection of vehicle, FGF1, or FGF19 (n=4-5/grroup one-way ANOVA F(2,3.46)=54.83 p=0.00241).

Next, we sought to compare the degree to which hypothalamic ERK1/2 signaling is induced following icv injection of FGF1 to that induced by icv FGF19, an endocrine member of the FGF family that transiently improves systemic glucose metabolism in diabetic rodents^10,14-17^. Prior work has shown that icv FGF19 induces ERK1/2 signaling in the hypothalamus and that improved glucose-tolerance in response to icv FGF19 requires functional ERK1/2 signaling^10^. Acutely, both FGF1 and FGF19 rapidly induce activation of ERK1/2 signaling in the MBH following icv injection^5,10^, however this effect persisted for 24h only after treatment with FGF1 (Figure 1E). Therefore, whereas both FGF1 and FGF19 rapidly activate ERK1/2 signaling in the hypothalamus following icv delivery, only FGF1 does so in a manner that is sustained for at least 24h.

### Diabetes Remission Induced by Central FGF1 Delivery Requires Functional Hypothalamic ERK1/2 Signaling

To determine whether sustained diabetes remission induced by icv FGF1 in diabetic Lep^*ob/ob*^ mice requires intact hypothalamic MAPK/ERK signaling, we co-administered the MAPK inhibitor U0126 with FGF1 and measured its impact on both hypothalamic ERK1/2 induction and on the duration of glucose lowering. We found that whereas activation of hypothalamic ERK1/2 signaling following icv administration of FGF1 (3 μg) was acutely blocked by pre-treatment with a single icv injection of U0126 (5 μg) (Figure 2A), this intervention did not prevent FGF1-induced reductions of food intake or body weight, nor did it block sustained lowering of the blood glucose level (Figures 2B–D). To explain this finding, we examined the duration of the effect of icv injection of U0126 (5 μg) on induction of hypothalamic ERK1/2 by icv administration of FGF1 (3 μg). We found that although this U0126 pre-treatment regimen was effective early on, it did not attenuate FGF1-induced activation of hypothalamic ERK1/2 at the 24h time point (Figure S1A).

**Figure 2.**
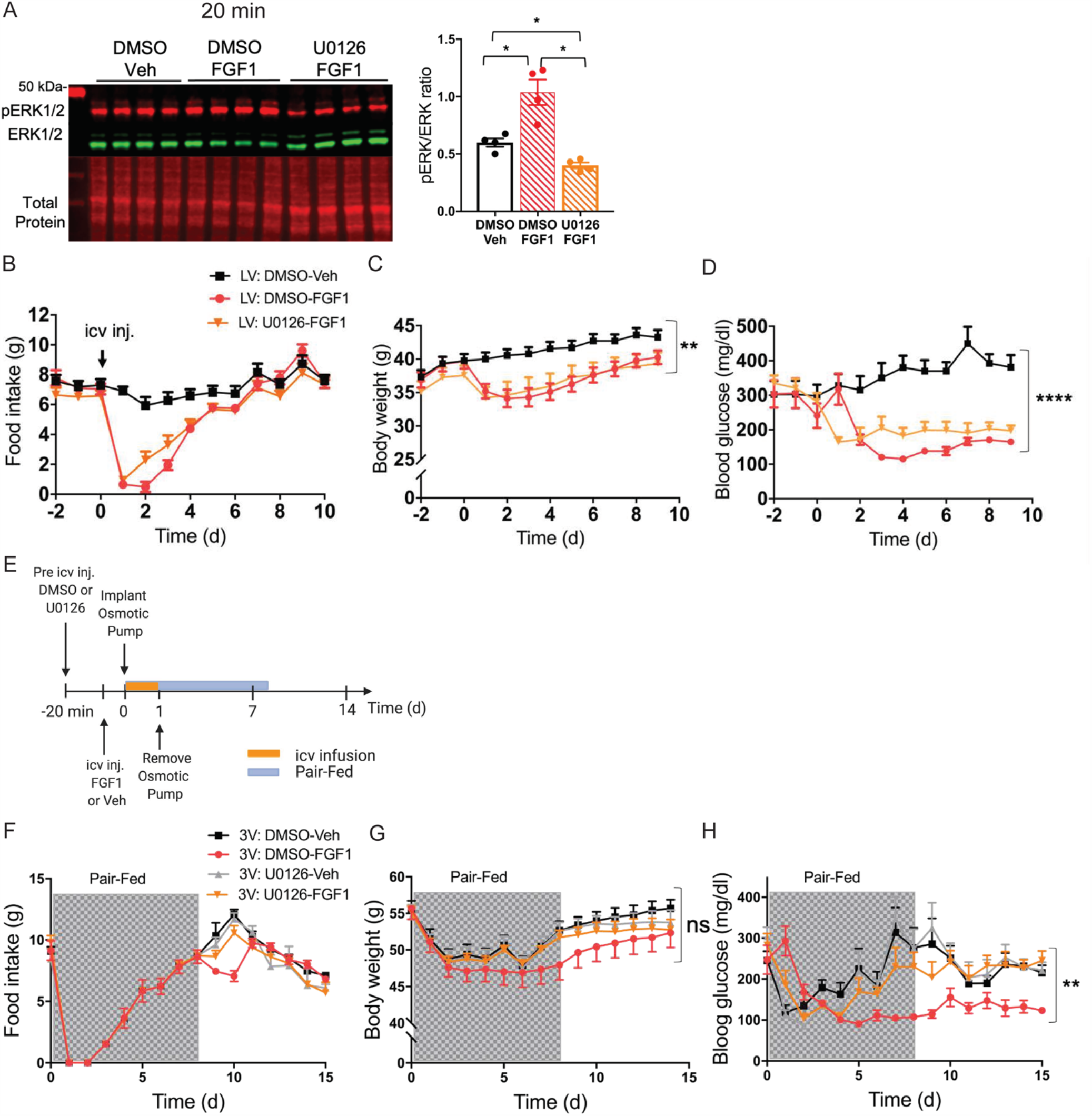
Prolonged MAPK signaling in the MBH is required for diabetes remission induced by central FGF1. **A)** Representative western blot (left panel) showing phosphorylated (red) and total ERK1/2 (green) and total protein (red), and data quantitation right panel) from hypothalamic punches from adult male C57Bl6J mice at 20 min after a injection into the lateral ventricle of the inhibitor of MAPK signaling U0126 (5 μg) or DMSO, followed by injection of either vehicle or FGF1 (n=5/group, one-way ANOVA F(2-5.27)= 20.692 p=0.00339). Effects of the same treatments on levels of **B)** food intake, **C)** body weight (n=7-8/group; repeated measures nparLD ANOVA.test statistic treatment =4.61 df=1.91 p=0.01) and **D)** blood glucose (n=7-8/group repeated measures nparLD ANOVA.test statistic treatment =27.96 df=1.8 p=7.36e-12) in Lep^*ob/ob*^ mice. **E)** Strategy for MAPK/ERK inhibition by continuous 3V infusion of U0126 or Vehicle DMSO for 24 hours followed by disconnection of osmotic pump and metabolic phenotyping image created with BioRender.com. Levels of **F)** food intake **G)** Body weight (n=5-8/group repeated measures nparLD ANOVA.test statistic treatment=0.80 df=2.70 p=0.47), and **H)** blood glucose (n=5-8/group repeated measures nparLD ANOVA.test statistic treatment =4.99 df=2.67 p=0.002) measured for 15 days post-treatment with icv injection of either saline vehicle or FGF1 (3ug) followed by a 24h continuous infusion of U0126 or DMSO vehicle into the third ventricle.

This observation prompted us to identify a route, dose, and duration of icv U0126 treatment capable of blocking FGF1-induced ERK1/2 activation for a full 24h period. Whereas neither repeated icv injection of U0126 (Figure S1B) nor continuous infusion of U0126 into the lateral ventricle (Figure S1C) were effective, we asked whether administration into the 3^rd^ ventricle would be more effective, owing to its close proximity to the MBH. Indeed, we found that continuous infusion of U0126 into the 3^rd^ ventricle for 24h (by osmotic mini-pump) following a single intra-3^rd^ ventricular U0126 injection was sufficient to block hypothalamic FGF1-induced ERK1/2 activation for a full 24h (Figure S1C).

We therefore employed this 24h 3^rd^ ventricular U0126 administration protocol to investigate the role played by MBH MAPK/ERK1/2 signaling in metabolic responses elicited by icv FGF1 injection (Figure 2E) in diabetic Lep^*ob/ob*^ mice. We found that whereas this intervention did not prevent initial reductions of food intake, body weight, or blood glucose induced by icv injection of FGF1 (3 µg) (Figures 2F– H), it did prevent sustained remission of diabetic hyperglycemia (Figure 2H). Because our previous work has shown that the initial glucose-lowering effect of icv FGF1 injection is driven primarily by transient anorexia^3,4^, a group of vehicle-treated Lep^*ob/ob*^ mice that were pair-fed to the intake of the group receiving FGF1 was included in this study. Data from this group demonstrates that in animals receiving icv vehicle treatment, pair-feeding to match the reduced food intake induced by icv FGF1 injection was sufficient to lower blood glucose levels initially. However, this glucose-lowering effect was not sustained irrespective of whether pair-feeding was accompanied by icv U0126 or vehicle administration (Figure 2E). Nevertheless, in mice that received both icv FGF1 injection and the 24h U0126 infusion, blood glucose levels returned to their elevated baseline value even before food intake and body weight had done so, whereas glucose levels remained low throughout the duration of the study in mice receiving FGF1 coupled with vehicle instead of U0126 (Figures 2E–H). Taken together, these results show that while hypothalamic ERK1/2 activation is not required for the acute, transient effects of icv FGF1 on levels of blood glucose, food intake, or body weight, it is required for sustained remission of diabetic hyperglycemia induced by the central action of FGF1.

### The FGF1 Mutant FGF1 R50E Mimics the Effect of Native FGF1 to Rapidly Induce ERK1/2 Activation in the Hypothalamus

As an alternative test of the hypothesis that diabetes remission induced by icv FGF1 injection requires sustained ERK1/2 activation in the hypothalamus, we repeated these studies in mice using FGF1 R50E, a FGF1 mutant in which arginine at the amino acid position 50 is substituted with glutamate^11^. In cell culture studies, FGF1 R50E retains the ability to bind to and activate FGFRs and induce transient activation of ERK1/2, but the effect is not sustained^11,12,18^. Consistent with these observations, we found that when given at the same dose (3 µg), activation of hypothalamic ERK1/2 by icv injection of FGF1 R50E was comparable to that induced by native FGF1 at the first 3 time points (20 min, 5h and 8h), but no effect was observed at either 14h or 24h after icv injection (Figure 3A). Therefore, FGF1 R50E-induced ERK1/2 activation in the hypothalamus is short-lived in comparison to the more durable response elicited by native FGF1 following icv injection.

**Figure 3.**
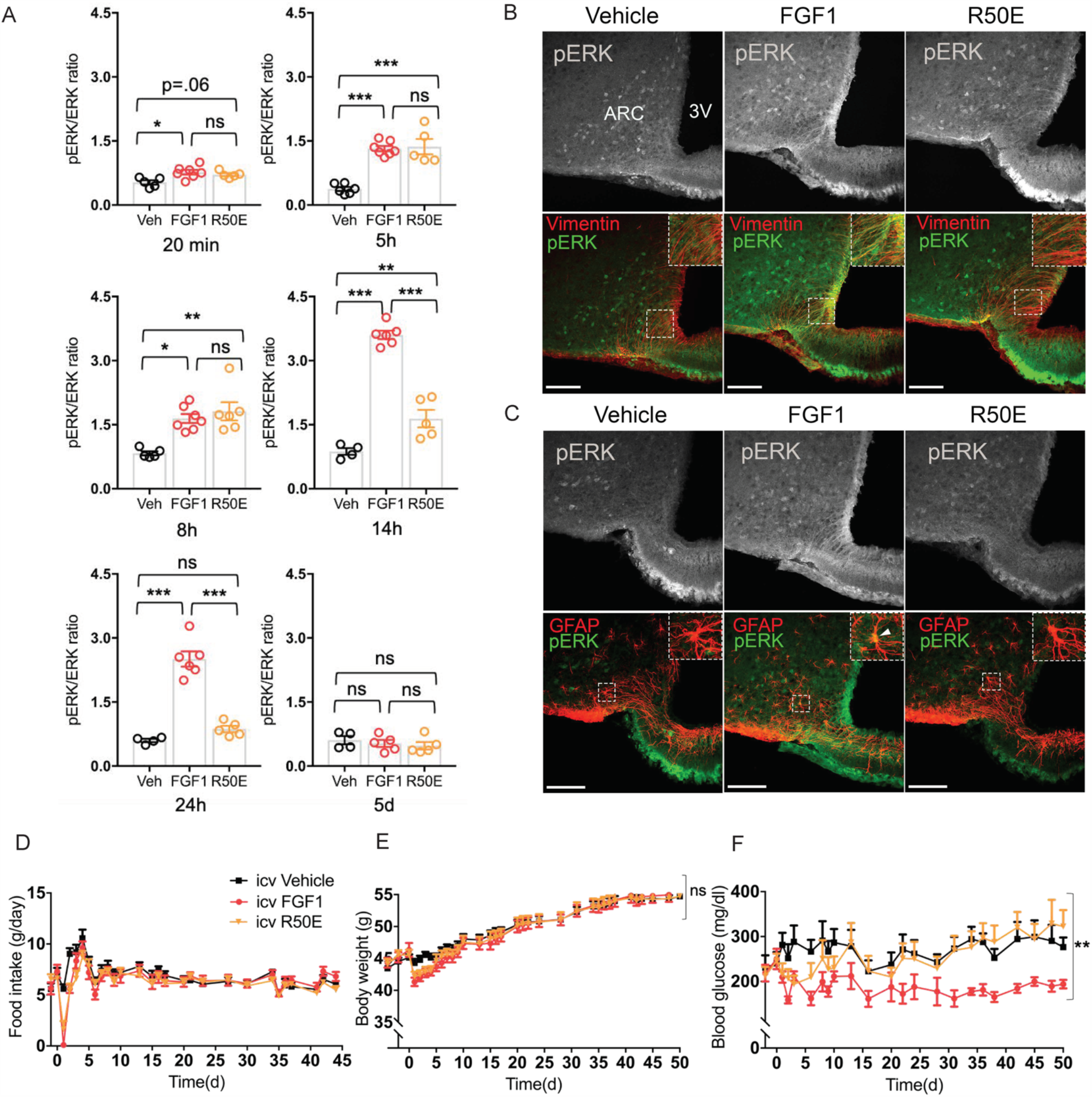
FGF1 Mutant Induces Only Transient ERK1/2 Activation in Hypothalamus and Fails to Induce Sustained Remission of Diabetic Hyperglycaemia. **A)** Quantitative western blot showing a time course of hypothalamic ERK1/2 phosphorylation after a single icv injection of either vehicle, FGF1 (3µg), or FGF1 R50E (3µg) at the following time points after injection: 20 min (n=5-7/group one-way ANOVA F(2,14)=6.1616 p=0.012); 5h (F(2,15)=31.334 p=4.4e-06) 8h (kruskal-wallis chi-squared=10.361 df=2 p=0.0056); 14h (n=4-6/group F(2,12)=101.94 p=2.9508) e-24h (F(2,12)=63.339 p=4.17e-07); or 5d (n=4-5/group F(2,11)=0.51 p=0.6) with Tukey’s post-hoc test ns=not significant, **P<0.01, ***P<0.001, ****P<0.0001. **B)** confocal images of the arcuate nucleus (ARC) and third ventricle (3V) showing pERK1/2 (grey) with merged pERK1/2 (green) and vimentin (red) and **C)** pERK1/2 (grey) with merged pERK1/2 (green) and glial fibrillary acidic protein (GFAP) (red) 14h after a single icv injection of either vehicle, FGF1(3µg) or FGF1 R50E (3µg). Scale bar 100 um and inserts representative double-labeled cells. Time course of the effect of the same 3 icv treatments on **D)** food intake, **E)** body weight (n=8-9/group repeated measures nparLD ANOVA.test statistic treatment=0.05 df=1.91 p=0.94), and **F)** plasma glucose levels (n=8-9/group repeated measures nparLD ANOVA.test statistic treatment=5.49 df=1.79 p=0.005) in Lep^*ob/ob*^ mice.

Based on previous work showing that tanycytes and astrocytes are especially responsive to the effect of icv FGF1 injection to induce sustained ERK1/2 signaling^5,7^, we next examined the response of these cell types following icv injection of FGF1 R50E. As anticipated, we observed that compared to icv injection of native FGF1, pERK1/2 induction in both tanycytes (Figure 3B) and astrocytes (Figure 3C) was markedly reduced at the 14h time point after icv injection of FGF1 R50E. These findings confirm that although ERK1/2 signaling is induced in tanycytes and astrocytes following icv administration of either FGF1 or FGF1 R50E, the response to the latter is short-lived compared to the highly durable response to native FGF1.

### FGF1 R50E does not Induce Sustained Remission of Diabetic Hyperglycemia

Having established that unlike native FGF1, the FGF1 R50E mutant does not induce sustained activation of ERK1/2 signaling in the hypothalamus following icv administration, we next sought to determine whether this blunted response corresponds to a reduced ability to induce sustained glucose lowering in diabetic Lep^*ob/ob*^ mice. We report that in diabetic Lep^*ob/ob*^ mice, icv injection of FGF1 R50E effectively recapitulated the acute reductions of food intake, body weight, and blood glucose induced by native FGF1 (Figures 3D–F). Whereas icv injection of native FGF1 induced remission of diabetic hyperglycemia that was sustained for at least 50 d, consistent with previous reports^3^, diabetic Lep^*ob/ob*^ mice treated icv with FGF1 R50E exhibited a complete relapse of hyperglycemia within 14 d of icv injection, coincident with recovery of baseline food intake and body weight (Figures 3D–F). Thus, whereas centrally administered FGF1 R50E mimics the acute but transient effects of native FGF1 on hypothalamic ERK1/2 activation, food intake, weight loss and blood glucose, neither the ERK1/2 activation nor the diabetes remission are sustained in the manner seen with icv FGF1 injection. This set of outcomes effectively replicates what is observed when icv FGF1 is co-administered with the MAPK inhibitor U0126 for 24h, and together these data offer direct evidence that the highly durable antidiabetic action of FGF1 in the brain is dependent on sustained hypothalamic ERK1/2 activation.

### Transcriptional Analysis of the Hypothalamic Response to Centrally administered FGF1 and FGF1 R50E at the Single-Cell Level

To characterize transcriptional differences elicited by sustained versus transient ERK1/2 activation engaged by FGF1, we leveraged a previously unpublished single-cell RNA-sequencing (scRNA-seq) dataset generated as part of a larger study investigating transcriptomic effects of icv injection of FGF1 in the MBH. That unpublished data was generated based on hypothalami from Lep^*ob/ob*^ mice icv injected with FGF1 R50E 5 days prior, and generated simultaneously with the hypothalamic scRNA-seq data from the Lep^*ob/ob*^ mice icv injected with either FGF1 or vehicle also 5 days prior (Figure 4A)^7^. This analysis revealed that in astrocytes, tanycytes, and oligodendrocyte lineage cells, the effect of icv FGF1 injection to induce differentially expressed genes (DEGs; False-discovery rate, FDR<0.05 and |log_2_ fold change|>0.25) was far greater than was observed following icv injection of the same dose of FGF1 R50E (Figure 4B). Nevertheless, correlation and ranking of differential gene expression by gene (see Methods) revealed that most genes upregulated by FGF1 were also upregulated by FGF1 R50E, but at much lower intensities (Figures 4C-D). From this, we infer that in the hypothalamus of Lep^*ob/ob*^ mice, transient and sustained ERK activation (induced by icv injection of FGF1 R50E and FGF1, respectively) induce the same gene expression programs but at differing magnitudes. This interpretation is further supported by geneset variation analysis of REACTOME pathways. We found that the majority of ‘ERK/MAPK’ associated pathways showed a significant change in activity, specifically in astrocytes, in response to treatment (Figure 4E). Using weighted gene co-expression network analysis (WGCNA)^19^ we further found that gene co-expression modules differed significantly (linear mixed-effects models; FDR<0.05; Supplementary Table 1) between FGF1 and FGF1 R50E treatment in astrocytes, but not other cell types. Specifically, modules ME11 and ME13 were significantly increased in astrocytes from mice receiving FGF1, but not FGF1 R50E, compared to icv vehicle treatment, whereas module ME5 was significantly reduced with FGF1 injection, but not with FGF1 R50E. Of interest is that module M13 is significantly enriched (p=0.026) for genes previously reported to be induced upon astrocyte-neuron interaction^20^. These observations support a model in which astrocyte activation resulting sustained FGF1-induced ERK1/2 signaling favors increased astrocyte-neuron interaction. Whether and how such an effect might induce sustained remission of diabetes is an important unanswered question.

**Figure 4.**
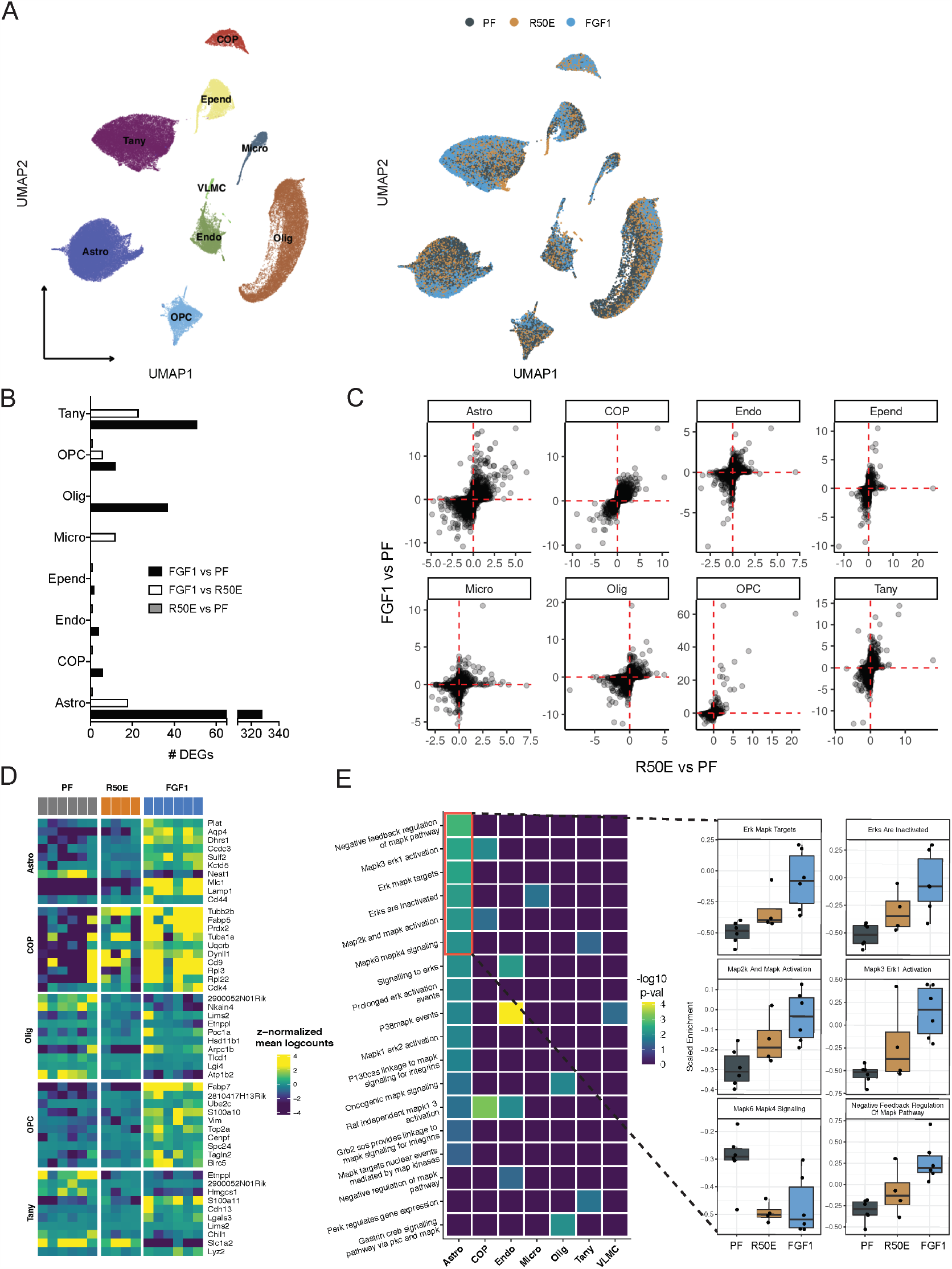
Hypothalamic Single-Cell Transcriptomics Reveal Transcriptional Differences between FGF1 and FGF1 R50E. **A)** Uniform manifold approximation and projection (UMAP) clustering of hypothalamic glia cells from diabetic Lep^*ob/ob*^ mice 5 days after a single icv injection of FGF1 or vehicle (GSE153551) or R50E. **B)** Number of differentially expressed genes (DEGs) identified through repeated downsampling (400 cells/group/cell type/iteration: 100 iterations). **C)** Correlation plot of DEGs ranked by |log2 fold change| multiplied with FDR. **D)** Top 10 most upregulated genes by FGF1 compared to pair-fed and FGF1 R50E in selected cell types. **E)** Geneset enrichment analysis of REACTOME terms related to ‘MAPK/ERK’ targets. Insert: scaled enrichment of top 6 enriched terms by treatment in astrocytes.

## Discussion

In the current work, we sought to determine the role played by increased hypothalamic MAPK/ERK signal transduction (measured by pERK1/2 western blot and immunostaining) in the sustained antidiabetic action elicited by icv injection of FGF1 in the Lep^*ob/ob*^ mouse model of T2D. Our findings demonstrate that *1)* activation of hypothalamic MAPK/ERK signaling is sustained for at least 24h following central administration of FGF1, and *2)* the ability of icv FGF1 injection to induce sustained normalization of diabetic hyperglycemia is dependent on this MAPK/ERK signaling response, but only if it is prolonged. In contrast, hypothalamic MAPK/ERK signaling does not appear to be required for transient effects on food intake, body weight, and blood glucose levels induced by icv FGF1 administration. These findings identify a key role for prolonged hypothalamic MAPK/ERK signaling in the sustained antidiabetic action of FGF1 action in the brain.

MAPK/ERK pathway activation is well documented following the binding of FGFRs by FGF1^9^, and a role for hypothalamic ERK signaling in the regulation of glucose homeostasis (in response to administration of leptin, insulin and FGF19) has been reported previously^10,21-23^. Based on these observations, along with our recent work showing that *1)* following icv FGF1 injection, robust induction of ERK1/2 signaling is observed in the hypothalamic arcuate nucleus-median eminence (ARC-ME)^5^, and *2)* FGF1 microinjection to this brain area recapitulates sustained glucose lowering observed following icv FGF1 injection^5^, we sought to investigate the role of hypothalamic MAPK/ERK signaling in FGF1-induced sustained glucose-lowering. To this end, we first characterized the time course of hypothalamic pERK1/2 induction following icv injection of FGF1 at a dose known to elicit sustained glucose lowering. Consistent with *in vitro* evidence that FGF1 induces both acute and sustained activation of ERK1/2 signaling^13^, we report that following a single icv FGF1 injection, hypothalamic pERK1/2 induction is observed within 20 min and persists for a full 24h.

Our next goal was to determine if this pattern of ERK1/2 activation is required for the sustained antidiabetic effect of icv FGF1, and two separate approaches were employed. First, we administered U0126, a potent and selective inhibitor of MAPK, as a single icv injection prior to icv administration of FGF1. Although this approach proved effective in terms of blocking FGF1-induced ERK1/2 activation in the short-term, MAPK blockade by icv infusion of U0126 over a full 24h period was required to block reversal of diabetic hyperglycemia in response to icv FGF1 injection. In contrast, this intervention had no impact on FGF1-induced anorexia, weight loss and associated transient glucose lowering, suggesting that unlike sustained glucose lowering, these responses to icv FGF1 are not dependent on ERK1/2 activation. Collectively, these data suggest that the sustained antidiabetic action of icv FGF1 involves mechanisms distinct from those involved in control of food intake, with the former, but not the latter, being dependent upon sustained ERK1/2 signaling in the hypothalamus.

Our second approach to testing this hypothesis took advantage of the fact that while the FGF1 R50E mutant activates FGFRs, it elicits on only transient activation of ERK1/2 in cultured cells^18^. Thus, we predicted that the response to icv injection of the FGF1 R50E mutant would mimic what is observed when the ability of icv FGF1 to induce sustained MAPK/ERK signaling is blocked by central administration of U0126. Indeed, we found that although FGF1 R50E induces hypothalamic ERK1/2 activation following icv injection, the effect is short-lived compared to that induced by icv FGF1 injection. Moreover, icv injection of FGF1 R50E fully recapitulated the effects of native FGF1 on food intake, body weight and associated transient amelioration of hyperglycemia, but it failed to induce sustained glucose lowering. Using two complementary approaches, therefore, our findings offer direct evidence that the MAPK/ERK signal transduction mechanism plays a key role in the effect of FGF1 to elicit sustained diabetes remission, while an unrelated signaling pathway mediates FGF1’s feeding effects.

Relevant to these observations is the mechanism engaged by FGF1-FGFR interaction that leads to sustained MAPK/ERK signal transduction. We suspect a role for integrins, a family of cell adhesion receptors that interact with FGFRs to modulate both receptor activity and the associated intracellular response^11,18,24^. The more durable and robust activation of intracellular signal transduction that results from FGFR-integrin co-activation is implicated in DNA synthesis, cell proliferation, cell migration, and other sustained cellular responses to growth factor signaling^11,25^. Integrins are also implicated in the control of insulin sensitivity in peripheral tissues in skeletal muscle and liver^26,27^, but not in the regulation of glucose homeostasis by the brain. As the FGF1 R50E mutant was previously characterized (based on work in cell culture) as being unable to activate FGFR1 without engaging integrin signaling, we favor a model in which the latter is induced in the hypothalamus by native FGF1, but not FGF1 R50E, and that this effect is required for sustained ERK1/2 activation. Investigation into the contribution made by brain integrin signaling to the antidiabetic effect of FGF1 is a priority for future work.

The interpretation of findings reported herein is also informed by recent work that has shed light on how the hypothalamus responds to FGF1. For example, intact hypothalamic melanocortin signaling (determined by interactions between Agrp and Pomc neurons in the ARC) appears to be required for sustained glucose lowering induced by icv FGF1 injection, but is dispensable for acute, transient effects of FGF1 on food intake, body weight, and glycemia^7^. In light of our current findings, this work suggests that diabetes remission induced by icv FGF1 involves a MAPK/ERK-dependent mechanism for enhanced melanocortin signaling. Whether this mechanism involves direct effects of FGF1 on Agrp and/or Pomc neurons or instead is mediated via an action on glial cells (*e*.*g*., tanycytes or astrocytes) warrants additional study, but the associated finding that FGF1 promotes cellular contacts between astrocytes and Agrp neurons supports the latter possibility^7^.

Also reminiscent of the current findings is our recent report that hypothalamic perineuronal nets (PNNs, defined as extracellular matrix specializations combined of proteoglycans and hyaluronan), which were recently shown to enmesh neurons in the ARC-ME^28^, are also targets for the glucose-lowering action of FGF1. Indeed, PNN formation increases rapidly in the ARC-ME following icv FGF1 injection and, in a rat model of T2D, disruption of this PNN response blocks the sustained antidiabetic action induced by icv FGF1 injection, while having no impact on the acute, transient effects of FGF1 on food intake, body weight, and glycemia^8^. These observations once again parallel those observed when hypothalamic MAPK/ERK signaling either is not induced or is blocked in a sustained manner. As such, they raise the interesting possibility that FGF1-induced stimulation of ARC-ME PNN formation – which likely involves FGF1 action on glial cells – depends upon MAPK/ERK signaling.

Additional insight into these possibilities is provided by comparing transcriptional changes occurring in the hypothalamus of Lep^*ob/ob*^ mice 5 days after icv injection of FGF1 *vs*. FGF1 R50E. While far more DEGs were induced by the former than the latter peptide, a closer examination reveals that the pattern of differential gene expression is in fact similar – what is different is the degree to which genes are induced. When the level of gene expression against known ERK1/2-mediated pathways was interrogated specifically in astrocytes, a similar pattern was detected. Namely, induction of genesets in the ERK1/2 pathway was more robust following icv FGF1 than FGF1 R50E when compared to pair-fed vehicle control mice, but the overall pattern of genesets was similar between the two. Using a different analytic approach, we further found that a co-expressed gene module detected in astrocytes following icv FGF1 injection, one that was not induced by injection of FGF1 R50E, is enriched for genes that are induced when astrocytes are co-cultured with neurons^20^. Although additional studies are needed to establish the role of astrocyte-neuron interaction in the hypothalamic response to FGF1, these data support a model in which sustained ERK1/2 activation initiates a gene co-expression program in astrocytes that allow them to interact with MBH neurons in ways that profoundly impact neurocircuitry involved in controlling glucose homeostasis.

We conclude that sustained diabetes remission induced by icv injection of FGF1 involves a robust and highly durable MAPK/ERK response in the hypothalamus, whereas the associated transient anorexia and loss of body weight induced by FGF1 does not. Although additional work is needed, convergent findings point to a role for prolonged MAPK/ERK signaling in hypothalamic glial cell types as primary targets for the glycemic benefit stemming from FGF1 action in the hypothalamus. Ongoing studies will continue to clarify the role of glia-neuron interactions in the ability of the brain to ameliorate hyperglycemia in a sustained manner in response to FGF1.

## Methods

### Resource Availability

#### Lead Contact

Further information and requests for resources and reagents should be directed to and will be fulfilled by the Lead Contact Jarrad M. Scarlett

#### Materials Availability

This study did not generate new unique reagents

#### Data and Code Availability

The datasets generated during this study are available at NCBI Gene Expression Omnibus (GEO) under Super Series number GSE153551 (https://www.ncbi.nlm.nih.gov/geo/query/acc.cgi?acc=GSE153551). Names, references, and version numbers of all software packages used are stated in the Methods section and in the Key Resource Table. Codes are available upon request.

### Experimental Model and Subject Details

#### Animals

Male, 8-week-old C57BL/6J (WT) and Lep^*ob/ob*^ (B6.Cg-Lep^ob^/J) mice were purchased from Jackson Laboratory (Bar Harbor, ME) and male, 8-week-old Lep^*ob/ob*^ (B6.V-Lep^ob^/JRj) were purchased from Janvier Labs, France. All animals were housed individually under specific pathogen-free conditions in a temperature control environment with either a 12 h: 12h or 14 h:10 h light: dark cycle, with 75-80% humidity and ad libitum access to water and standard laboratory chow (LabDiet, St Louis, MO) or Altromin 1310 chow (Brogaarden, Denmark). All animal procedures were performed according to the National Institutes of Health Guide for the Care and Use of Laboratory Animals and approved by the Institutional Animal Care and Use Committee at the University of Washington or performed with approved protocols from The Danish Animal Experiments Inspectorate permit number 2014-15-0201-00181 and the University of Copenhagen project number P16-122.

## Method Details

### Surgery

Lateral ventricle (LV) and third ventricle (3V) cannulation (8IC315GAS5SC, 26-ga, Plastics One, Roanoke, VA) were performed under isoflurane anesthesia using the following stereotaxic coordinates for mice: LV: −0.7 mm posterior to bregma; 1.3 mm lateral, and 1.3 mm below the skull surface and 3V: −1.8 mm posterior to bregma; mid-line and −4.3 mm below the skull surface. Animals received buprenorphine hydrochloride (Reckitt Benckiser Pharmaceuticals Inc., Richmond, VA) for pain relief and were allowed to recover for one week prior to the study.

### Icv injections

Mean blood glucose and body weight values were matched between groups before the icv injections. Animals received a single icv injection via the LV of either saline vehicle, recombinant mouse FGF1 (FGF1; Prospect-Tany TechnoGene Ltd., East Brunswick, NJ) dissolved in sterile water at a concentration of 1.5 μg/µl, recombinant human FGF1 (a generous gift from Novo Nordisk) dissolved in PBS pH 6.8, human FGF1 R50E (a generous gift from Novo Nordisk) dissolved in 20 mM Tris pH 8.0, 0.5 M NaCl or human FGF19 (hFGF19; Phoenix Pharmaceuticals, Inc., Burlingame, CA) dissolved in 0.9% normal saline at a concentration of 1.5 μg/µl. and injected using a 33-gauge needle (8IC315IAS5SC, 33 ga, Plastics One, Roanoke, VA) extending 0.8 mm beyond the tip of the icv cannula over 60 s for a final volume of 2 μl^3^.

To determine if the effect of icv FGF1 administration to induce sustained remission of hyperglycemia in Lep^*ob/ob*^ mice depends on sustained MAPK/ERK signaling, we studied four groups of diabetic *Lep*^*ob/ob*^ mice. Two groups received a single icv injection of FGF1, while the other two received icv vehicle injected into the 3V using a 33-gauge needle (8IC315IAS5SC, 33 ga, Plastics One, Roanoke, VA) extending 1mm beyond the tip of the icv cannula. Each group also received a pre injection and an icv infusion of either the selective MAPK inhibitor U0126 (662005; Millipore Sigma, St. Louis, MO) dissolved in DMSO at a concentration of 30mM or DMSO vehicle 1 μl per hour into the 3V via an micro-osmotic pump (model 1003D; Alzet, Durect Corporation, Cupertino, CA) connected to the 3V cannula for 24 hours.

### Quantitative western blotting

Hypothalamic punches (3 mm) were collected after indicated treatment. Dissection of a single coronal section (1.5 mm) between the rostral and caudal Circle of Willis using an ice-cold brain matrix followed by a 3 mm biopunch (Harris Uni-Core, Ted Pella Inc, Redding, CA) of the MBH was flash-frozen in liquid nitrogen and stored at −80 °C until further processing. Hypothalamic punches were homogenized by sonication in lysis buffer and centrifuged at 10,000 g for 15 minutes supernatant was collected and assayed for protein concentration using a BAC assay. Lysates were mixed in the Licor protein sample loading buffer and heated to 100 °C for 5 minutes. Samples were run on a 10% Bis-Tris criterion XT gel (Bio-Rad Laboratory Inc., Hercules, CA) at 200 volts for 25 minutes electrophoretic transferred at 100 V for 45 minutes. Loaded protein concentrations were determined based on primary antibodies 1:1000 rabbit anti-pERK1/2 antibody (#4370; Cell Signaling Technology, Danvers, MA), mouse anti-ERK1/2 (#9107; Cell Signaling Technology, Danvers, MA), secondary IRDye 680RD (#926-68073 Li-cor, Lincoln, NE), IRDye 800CW (#92632212; Li-cor, Lincoln, NE) and total protein Revert 700 Total Protein Stain Kit (#926-11011; Li-cor, Lincoln, NE) combined linear range.

### Immunofluorescence

Activation of the MAPK/ERK pathway at 14 hours post icv injection pERK1/2 was detected by immunohistochemistry in overnight-fasted C57Bl6J mice. Fourteen hours following icv injection of either vehicle, recombinant FGF1 (3 ug), or FGF1 R50E (3 ug), mice were anesthetized with ketamine and xylazine and perfused with PBS followed by 4% paraformaldehyde in 0.1 mol/L PBS, after which brains were removed. Brain were placed in sucrose overnight. Cryostat sectioned (30-um thick) free-floating sections were permeabilized with 0.4% PBS-T overnight Wash in 0.4% PBS-T incubate in pERK1/2 antibody, Host: Rabbit, (#4370;Cell Signaling Technology, Danvers, MA) 1:1000 in (3%BSA+0.4%TritonX-100+ 0.2%Normal Donkey Serum in PBS-azide) for 24h at 4C Wash sections 5× 10min with 0.4% PBS-T Incubate in Secondary Antibody (DαR 594-A21207) 1:1000 in (3%BSA+0.4%TritonX-100+ 0.2%Normal Donkey Serum in PBS azide) for overnight in 4 C Incubate in DAPI 1:10,000 in PBS for 10min at room temperature and wash tissue 3× 10min with PBS followed by mount, dry and coverslip with PVA mounting media. Immunofluorescence images were captured using a Leica SP8X Scanning Confocal microscope (Leica Microsystems, Buffalo Grove, IL) with an HC FLUOTAR L 25X/0.95 W objective.

### Generation of single-cell suspensions

Diabetic Lep^*ob/ob*^ mice received an icv injection of either FGF1 (n=6), FGF1 R50E (n=4) or vehicle (n=6). Mice were euthanized 5 days later, and brains were extracted between 9-12 AM, cooled in ice-cold DMEM/F12 media (Gibco, Thermo Fisher Scientific) for 5 minutes. Brains were then placed ventral surface up into a chilled stainless steel brain matrix and a single coronal section (1.5 mm thick) block was obtained between the rostral and the caudal ends of the Circle of Willis, followed by a triangular section of the MBH using a scalpel. Dissected tissue was digested using the Neural Tissue Dissociation Kit from Miltenyi Biotec (Bergisch Gladbach, Germany) with manual dissociation with the following modifications: sections were immediately placed in preheated (37°C) enzyme mix 1 (1425 ml Buffer X + 37.5 ml Enzyme E) and incubated in closed tubes for 15 minutes at 37 °C under slow, continuous rotation. Then enzyme mix 2 (15 ml Buffer Y + 7.5 ml Enzyme A) was added followed by mechanically dissociated using a wide-tipped, fire-polished Pasteur pipette by pipetting up and down 10 times slowly. After a 10-minutes incubation under slow, continuous rotation, using a series of 2 fire-polished Pasteur pipettes with incrementally smaller openings the tissue was gently dissociated until no visible pieces were observed. Cell suspensions were centrifuged at 300xg for 10 minutes at room temperature (RT), and resuspended in 500 ml Hanks’ Balanced Salt Solution (HBSS) (Gibco, Thermo Fisher Scientific) and 0.04% (w/v) bovine serum albumin (BSA) (Sigma-Aldrich). Following an additional wash (centrifugation at 300xg for 5 minutes and re-suspension in 100 µl HBSS+0.04% BSA) cells were filtered through a 40 mm mesh and 50 ml HBSS+0.04% BSA was added through the filter and transferred to clean tubes. The cell suspension was centrifuged at 300xg for 5 minutes and resuspended in 50-150 ml HBSS+0.04% BSA. Cell concentrations were estimated on a NucleoCounter NC-3000 (Chemometec, Denmark) and kept on ice until single-cell encapsulation.

### Single-cell RNA sequencing

Single-cell cDNA libraries were generated using the 10x Genomics (USA) Chromium single-cell controller and the 3’ v2 Reagent Kit according to manufacturer’s protocol. Single-cell libraries were sequenced on a NextSeq 500 platform with 3 samples on one flow cell to obtain 100 and 32-bp paired end reads using the following read length: read 1, 26 cycles, read 2, 98 cycles and i7 index, 8 cycles.

### Single-cell RNA-sequencing data processing

Cell Ranger version 1.2 (10x Genomics, USA) was used to de-multiplex and quantify unique molecular identifiers (UMI). A count matrix was generated for each sample with default parameters and genes were mapped to the mouse reference genome GRCm38 (mm10, part of the Cell Ranger software package; for the scRNA-seq dataset) or an adapted reference from the mouse reference genome including introns (for the snRNA-seq dataset), following the steps outlined on the 10x Genomics website (https://support.10xgenomics.com/single-cell-geneexpression/software/pipelines/latest/advanced/references#premrna). Barcodes were filtered to include those with a total UMI count >10% of the 99th percentile of the expected recovered cells. Next, Scrublet^29^ was run with default parameters to identify likely doublets which were subsequently removed from the expression matrix. Further processing was performed using the Seurat^30,31^ R package (version 2.3). Cells in which <400 or >4,000 expressed genes were detected, as well any cell with >20% mitochondrial and/or ribosomal transcripts were discarded. Counts were normalized using the SCtransform() function from Seurat. Variable genes were identified using the FindVariableGenes() function with default settings. Unwanted sources of variation, e.g. number of UMIs and ribosomal and mitochondrial percentage, were regressed out using the ScaleData() function. Principal components were identified with variable genes using the RunPCA() function. The PCs for clustering were identified using the maxLikGlobalDimEst function from the intrinsicDimension^32^ package which is described as best practice^33^. These 10 PCs were used to identify clusters and with the FindClusters() function at a resolution of 0.2. All neuronal lineage cells were removed from this dataset, as the quality was poor due to mouse age (myelination/neuron fragility).

This dataset was then merged with the previously published part of this dataset comprising of hypothalamic cells from vehicle and FGF1 treated mice. As cells merged into the expected clusters, no integration methods were required. Cluster labels from the previous dataset were used to annotate the clusters found in the merged dataset.

The final step was to further remove any artifacts/doublets, which had been missed in the initial round of processing. Each cell type underwent a round of iterative clustering as described above. Within each cell type, any cluster in which 50% of cells originated from a single sample was removed. The FindMarkers() function was used to identify markers of each subcluster, which were removed if enriched for marker genes of a different cell type.

Finally, for each cell, the silhouette index, a measure quantifying the similarity of a given cell to its assigned cluster compared to other clusters, was computed, and subsequently in order to remove cells that could not confidently be assigned to a single cluster, cells with a negative silhouette index were discarded.

### Single-cell RNA-sequencing analyses

#### Cell Type Abundance

Cell type abundance per individual sample was calculated by summing the total number of cells within each defined cell cluster and then dividing by the total number of cells collected per sample. Shifts in cell type abundance across conditions were tested using a mixed effects linear model with a random effect for samples to account for the hierarchical structure of the data.

#### Differential State analysis

Using the Muscat package() pre-defined clusters were split by sample requiring at least 10 cells per sample-cluster combination^34^. Raw counts were summed across each gene-sample-cluster combination to generate a single column per sample-cluster. This matrix was then fed directly into edgeR() for standard processing^35^. A gene was considered differentially expressed if the adjusted local p-value (adjusted within a cluster) was less than 0.05.

#### Correlation Analysis

Following DGE analysis, each gene was assigned a rank within each cluster treatment combination based on its differential expression (-log10(p-value) * log2FoldChange). The transcriptomic response to the multiple treatment paradigms were then compared to one another by calculating the spearman’s correlation of gene rankings.

#### GSVA analysis

Gene set variation analysis was performed using the R package ‘GSVA’ function with default parameter s(method=“gsva”)^36^. GSVA implements a non-parametric unsupervised method for gene set enrichment that transforms the data from a gene by sample matrix to a gene-set by sample matrix, allowing for individual sample gene-set enrichment testing. As input to GSVA, we calculated the pseudobulk profiles from each sample cluster combination which were then normalized with the variance stabilizing transformation function from DESeq2. All gene sets from the REACTOME database were tested for enrichment. The association between gene set enrichment and treatment status was assessed using linear regression. A linear model with the normalized enrichment score as dependent and treatment group as independent variable was constructed for each gene set. P-values were adjusted for multiple testing using the FDR correction.

#### WGCNA and adjacency matrix calculation

The WGCNA package (version 1.66) (cite number 17) implemented in R was used^37^. Pearson correlation coefficient was used for the single-cell data and signed network parameter was used to compute the gene-gene adjacency matrix. Powers corresponding to the top 95th percentile of network connectivity or above were discarded and the lowest soft threshold power between 1 and 30 to achieve a scale-free topology R-squared fit of 0.8 was selected.

#### WGCNA hierarchical clustering

The topological overlap matrices were converted to distance matrices, and the hclust() function was used with the complete method to cluster genes hierarchically. The cutreeDynamic() function was used with a deepSplit parameter of 2 and the pamStage parameter set to FALSE to carve the dendrogram into modules with at least 15 genes. The moduleEigengenes() function was used to compute module eigengenes, the vector of cell embeddings on the first PC of each module’s expression submatrix. The mergeCloseModules() function was used to merge modules, using a cutHeight of ≤0.2, corresponding to a Pearson correlation between module eigengenes of ≥0.8. The module eigengenes and expression matrices were used with the signedKME() function to compute gene–module Pearson correlations, or kMEs, a measure of how close each gene is to each module. Module treatment associations were tested using a linear mixed-effects model with the lme4 R package^38^ (function lmer()), using a random effect for each sample. Significance values are FDR corrected to account for multiple testing.

#### Astrocyte Enrichment

To characterize WGCNA modules found enriched in FGF1 treated astrocytes when compared to FGF1-R50E and vehicle treated mice, a hypergeometric enrichment test was performed using markers of astrocytes in middle cerebral artery occlusion (MCAO)/lipopolysaccharide (LPS) mice or astrocytes co-cultured with neurons. Prior to the analysis, the (GSE35338) ^39^ and (E-MTAB-5514) ^20^ datasets were downloaded and DEG analysis was performed. Genesets were constructed by identifying any gene with a log_2_ fold-change>2 and FDR<0.05 (omitted for the GSE35338^39^ dataset) in any of the conditions described. Any genes shared between LPS and MCAO at Day 1 were assigned to the PAN-reactive geneset. Furthermore, any gene found to overlap between LPS and MCAO was removed from the geneset. All p-values were FDR-corrected using sBH correction.

### Statistical Analysis

Data from individual experiment are shown as dot plots representing data from individual animals and bar graphs representing average ±SEM. Statistical analyses were performed using R ^40^. Student’s t-test was used to compare the means in two groups and one-way ANOVA to compare three groups assumptions of test were satisfied. For data that was collected repeatedly on each individual animal overtime we employed robust non-parametric methodology statistical procedures to enable accurate and reliable analysis of longitudinal measurements with minimal conditions and have competitive performance for small sample size when (semi)parametric assumptions were are not satisfied. Longitudinal data were analyzed using a linear mixed model including fixed effect of treatment and day and random effects of animal. Linear mixed models were conducted with the R statistical package “nlme”^41^, or the equivalent nonparametric test using the R statistical package “nparLD” ^42^ and followed up by post hoc analysis using nparcomp^43^. Probability values less than 0.05 were considered statistically significant.

## Acknowledgements

The authors are grateful for the technical assistance provided Helle Kinggaard Lilja-Fischer, Cecilia Ratner, Birgitte Holst, and the Single-cell Omics Platform at the Novo Nordisk Foundation Center for Basic Metabolic Research, University of Copenhagen. The authors thank Nathanial Peters at the University of Washington Keck Imaging Center for technical assistance and the National Institutes of Health (S10-OD-016240) for support to the W.M. Keck Foundation Center for Advanced Studies in Neural Signaling. JMB was supported by National Heart, Lung, and Blood Institute T32 training grant HL-007312 and the Diabetes Research Center Samuel and Althea Stroum Endowed Graduate Fellowship and HW was supported by the NIDDK-funded Diabetes, Obesity and Metabolism Training Grant (T32 DK007247) at the University of Washington. This work was supported by NIH-NIDDK grants: K08DK114474 (JMS), R01DK101997 (MWS), R01DK089056 (GJM), R01DK083042 (GJM and MWS) and an American Diabetes Association Innovative Basic Science Award (ADA 1-19-IBS-192; GJM). This work was also supported by the NIH-NIDDK–funded Nutrition Obesity Research Center (NORC; P30DK035816) and the Diabetes Research Center (DRC; P30DK017047) at the University of Washington. Additional funding to support these studies was provided to JMS by the UW Royalty Research Fund (RRF; A139339) and to THP by the Lundbeck Foundation (Grant number R190-2014-3904). Funding was also provided to MWS by Novo Nordisk (CMS-431104) and to MAB by the Novo Nordisk Foundation (NNF17OC0024328) and the Novo Nordisk Foundation Center for Basic Metabolic Research, which is an independent research center at the University of Copenhagen partially funded by an unrestricted donation from the Novo Nordisk Foundation (NNF10CC1016515).

## Author Contributions

JMB, MWS, and JMS conceived the project. JMB, MAB, THP, MWS, and JMS designed the experiments. JMB, MAB, DMR, BAP, DW, HW, MEM, NER and JMS, performed the experiments. JMB, MAB, DMR, BAP, XZ, PZ, AS, GJM, THP, MWS, and JMS acquired, analyzed, and interpreted the data. JMB, MAB, DMR, MWS, and JMS drafted and revised the manuscript. All authors approved the final version of the manuscript. JMS is the guarantor of this work and, as such, had full access to all of the data in the study and takes responsibility for the integrity of the data and the accuracy of the data analysis.

