## Supplemental Figure 1 for "Role of hypothalamic MAPK/ERK signaling in diabetes remission induced by the central action of fibroblast growth factor 1"

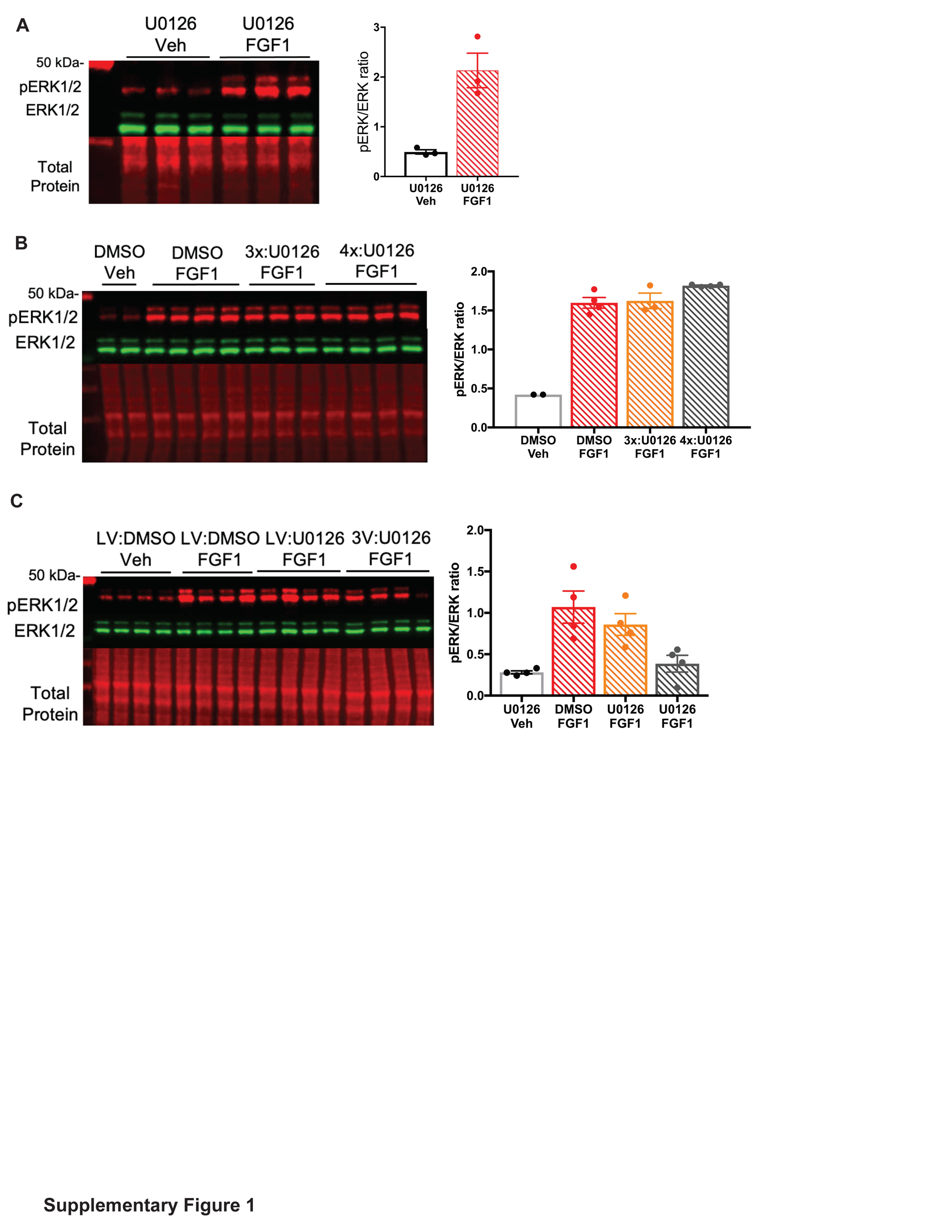

**Supplementary Figure 1- Blockade of Prolonged MAPK/ERK signaling in the MBH induced by central FGF1. A)** Representative western blot (left panel) and quantitative comparison of phosphorylated (red) and total ERK1/2 (green) and total protein (red) (right panel) from hypothalamic punches from adult male C57Bl6J mice 24h after a icv injection of an inhibitor of MAPK signaling U0126 (5 μg) followed by vehicle or FGF1 into the lateral ventricle and **B)** 24h after 3 and 4 repeated injections of U0126 (5 μg) 3h apart after a single icv injection of either vehicle or FGF1. **C)** a single icv injection of FGF1 into the lateral ventricle (LV) or 3rd ventricle (3V) followed by continuous infusion via osmotic pump of U0126 or DMSO for 24h.
